# The household resistome – frequency of beta-lactamases, class 1 integron and antibiotic resistant bacteria in the domestic environment

**DOI:** 10.1101/2020.06.04.135590

**Authors:** Laura Schages, Ralf Lucassen, Florian Wichern, Rainer Kalscheuer, Dirk Bockmühl

## Abstract

The widespread of antibiotic resistance (ABR) among bacteria has become a global health concern for humans, animals and the environment. In this respect, beta-lactams and colistin are of particular interest due to the emergence of multidrug-resistant gram-negative bacteria. Households provide a habitat for bacteria originating from humans, animals, foods, contaminated clothes or other sources in which detergents and biocides are frequently used. Thus, bacteria carrying antibiotic resistance genes (ARGs) might be introduced into private households and may consequently be also released from households to the environment via domestic wastewater. Since data on ABR in the domestic environment is limited, this study aimed to determine the abundance and correlation of beta-lactamase, mobile colistin resistance and class 1 integron genes and to characterize phenotypic resistant strains in private households in Germany. Additionally, the persistence of ABR bacteria to laundering and automated dishwashing was assessed. Shower drains, washing machines and dishwashers were sampled and analyzed using quantitative real-time PCR. Resistant strains were isolated, followed by identification and antibiotic susceptibility testing using VITEK 2. The results show a significantly higher occurrence of ARGs in shower drains compared to washing machines and dishwashers. Several beta-lactamase genes co-occurred and resistance of bacterial isolates correlated positively with genotypic resistance. Laundering and automated dishwashing reduced ABR bacteria significantly and the efficacy increased with increasing duration and temperature. Overall, the domestic environment seems to represent a potential reservoir of beta-lactamase genes and beta-lactam resistant bacteria with shower drains as the dominant source of ABR.

**Importance:** The abundance of ABR bacteria and ARGs is steadily increasing and has been comprehensively analyzed in natural environments, animals, foods or wastewater treatment plants. Despite of their connection to these environments, private households seem to be neglected. Therefore, the present study investigated shower drains, washing machines and dishwashers as possible sites of ARGs and ABR bacteria. The analysis of the domestic environment as a potential reservoir of resistant bacteria is crucial to determine whether households contribute to the spread of ABR or are a habitat where resistant bacteria from the environment, humans, food or water accumulate.

## Introduction

Antibiotics are extensively used in human and veterinary medicine, leading to the increase of antibiotic resistance (ABR) among bacteria and thus making it to a global health concern for humans, animals and the environment (1, 2). Besides overuse and misuse of antibiotics, the decreasing number of new drugs developed enhances the ABR crisis (3). Infections caused by multidrug-resistant (MDR) gram-negative bacteria such as *Enterobacteriaceae* or *Pseudomonas aeruginosa* occur worldwide (3–5). In this respect, resistance to beta-lactams and colistin is of great concern since these drugs are considered to be critically important antimicrobials for human medicine (1). Colistin is often used as an antibiotic of last resort for infections with MDR gram-negative bacteria (6, 7). Therefore, the increasing number and diversity of beta-lactamase (*bla*) genes (8) and mobile colistin resistance (*mcr*) genes (9) in pathogenic bacteria dramatically limit the treatment options with available antibiotics (10, 11). *Bla* and *mcr* genes are often located on mobile genetic elements (6, 12) or integrons (13, 14), enabling their horizontal and vertical transfer between bacteria. Class 1 integrons (*intI1*) have been identified as a proxy for pollution with heavy metals, antibiotics or personal care products and have already been used as a marker for the dissemination of antibiotic resistance genes (ARGs) (15, 16) since they are frequently connected with carrying multiple ARGs (17, 18).

ABR bacteria and ARGs have been comprehensively analyzed in humans, animals, food, soil, water and wastewater (19–25) while the domestic environment seems to be neglected. However, it seems reasonable that bacteria carrying ARGs might be introduced into private households by contaminated clothes, skin, food stuff or other sources and may consequently be also released from households to the environment via domestic wastewater of dishwashers, washing machines and drains. Since domestic wastewater must be considered as an important component of wastewater, households could also play a role in the dissemination of ABR bacteria and ARGs. However, data on the domestic environment is limited (16, 26, 27) although ABR bacteria in domestic appliances such as washing machines or dishwashers might pose a potential health risk to inhabitants, especially due to cross-contaminations of contaminated laundry or dishes (28–31). In an earlier study the occurrence of *bla* genes in domestic appliances was qualitatively analyzed and it was found that *bla* genes occurred in 79% of the washing machines and in 96% of the dishwashers, with ampC- and OXA-type genes dominating (26). Although the impact of domestic laundering was assessed in this study, the effect of automated dishwashing on ABR bacteria has not been determined so far. Based on these observations the present study aimed at quantifying the abundance of *bla, mcr* and *intI1* genes in washing machines, dishwashers and shower drains. We hypothesized that (i) ARGs and ABR bacteria are strongly abundant in households, that (ii) *bla* genes positively correlate with each other in households and that (iii) the reduction of ABR bacteria increases with higher temperatures and longer washing periods during automated dishwashing as well as laundering.

## Material and methods

### Sampling and sample preparation

Samples of shower drains, dishwasher sumps and sieves, detergent trays and rubber door seals of washing machines were taken from 54 households in North-Rhine-Westphalia, Germany between September 2018 and June 2019. All samples were taken in households without home care, from different apartment buildings or single family houses. Samples were collected by probing the surfaces of the inner tubing of shower drains (SD), the sumps and sieves of dishwashers (DW) or the detergent trays and rubber door seals of washing machines (WM) by swab method in triplicate.

Samples were stored immediately at 4 °C and prepared within 24 h. Samples collected by swab method were suspended in 1000 µL of sterile 0.9 % sodium chloride (three swabs were taken and pooled). After centrifugation at 4,800 rpm/8 °C/15 min the supernatant was discarded and the resulting pellet was re-suspended in 500 µL of sterile 0.9 % sodium chloride (16). The suspended samples were used for further analyses.

### DNA Extraction

For purification of total DNA, the Fast DNA Spin Kit for Soil (MP Bio, Santa Ana, CA, USA) was used according to the manufacturer’s instructions with the following adjustments: Instead of 500 mg solid material, 250 µL of suspended sample were applied to the lysing matrix tube. Samples were homogenized twice in the FastPrep-24™ instrument for 60 sec at 6.0 m/s. All samples were washed twice using the SEWS-M solution of the kit. After DNA-extraction real-time quantitative PCR (qPCR) was performed for the detection of resistance genes.

### Detection of *bla, mcr* and *intI1* genes

To detect *bla* and *mcr* genes, qPCR was performed as described by Rehberg et al. (2017). All sequences of the used oligonucleotides (custom synthesized by Biolegio, Nijmegen, Netherlands) are shown in table 1. The genes *bla*_ACT_ and *bla*_MIR_ will be referred to as *bla*_ACT/MIR_, since the oligonucleotides targeted both. The prepared DNA was amplified using the HotStarTaq^®^ Master Mix Kit (Qiagen, Hilden, Germany) and a mix of oligonucleotides and DNA probes labelled with different dyes. The PCR mix contained 10 µL of a 2x HotStarTaq^®^ Mastermix, 7 µL of RNase-free water, 1 µL of oligonucleotide mix (oligonucleotide: 4 µM; probe: 2 µM and MgCl_2_: 50 µM) and 2 µL of the DNA template comprising a final volume of 20 µL. The PCR was performed on a LightCycler 480 (Roche Life Sciences, Mannheim, Germany) using the following conditions: 95 °C for 15 min, 45 cycles of denaturation (95 °C; 10 s), annealing (60 °C; 20 s), elongation (72 °C; 10 s) and a final cycle at 30 °C for 30 s. A sample was considered positive if it reached the threshold before cycle 40 or if the copies/µL determined by standard curve were greater than five copies/µL. For determination of 16S ribosomal DNA (16S rDNA) and *intI1* genes, qPCR was performed according to Lucassen et al. (2019), primers are shown in table 2.

**Table 1.**
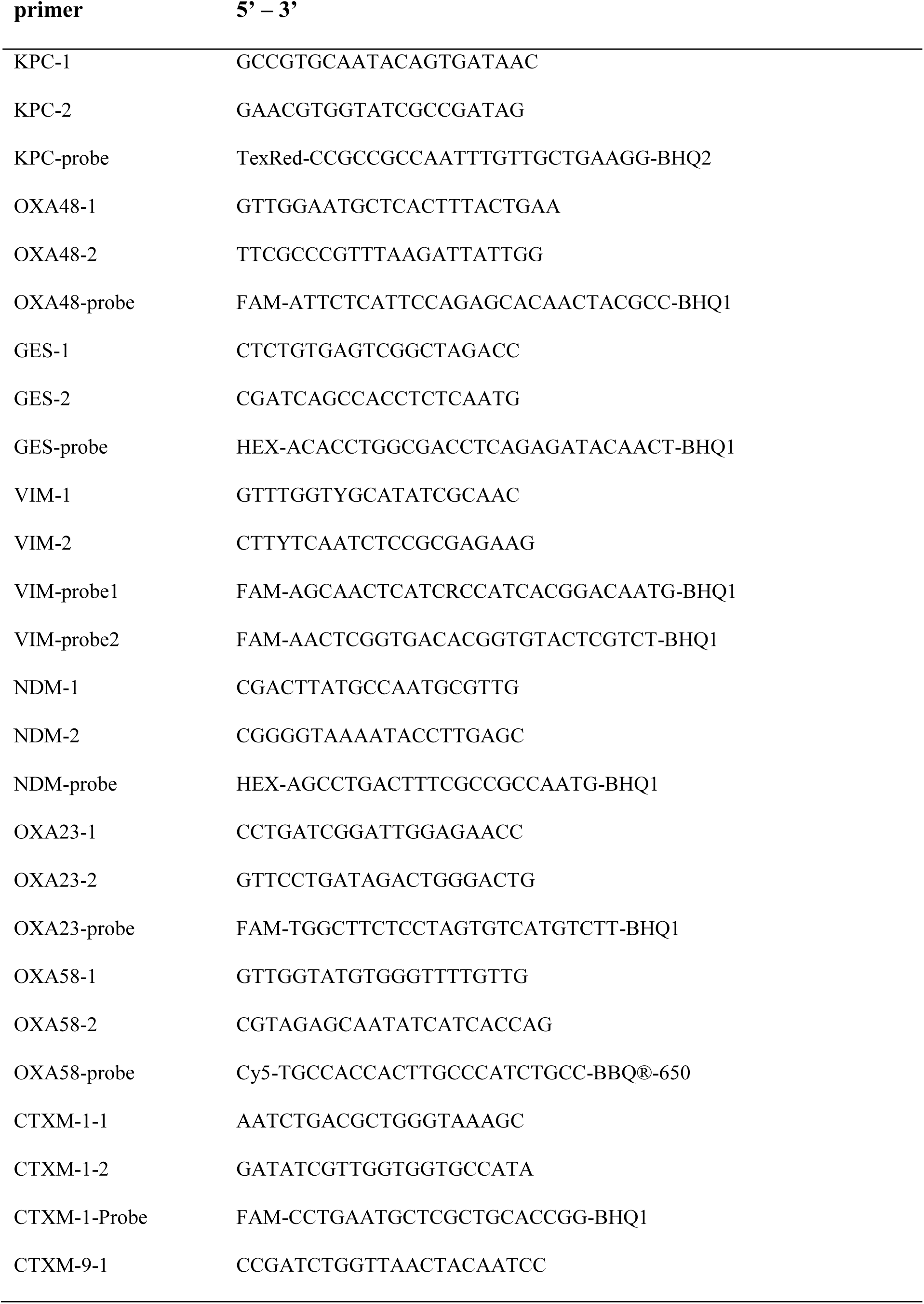
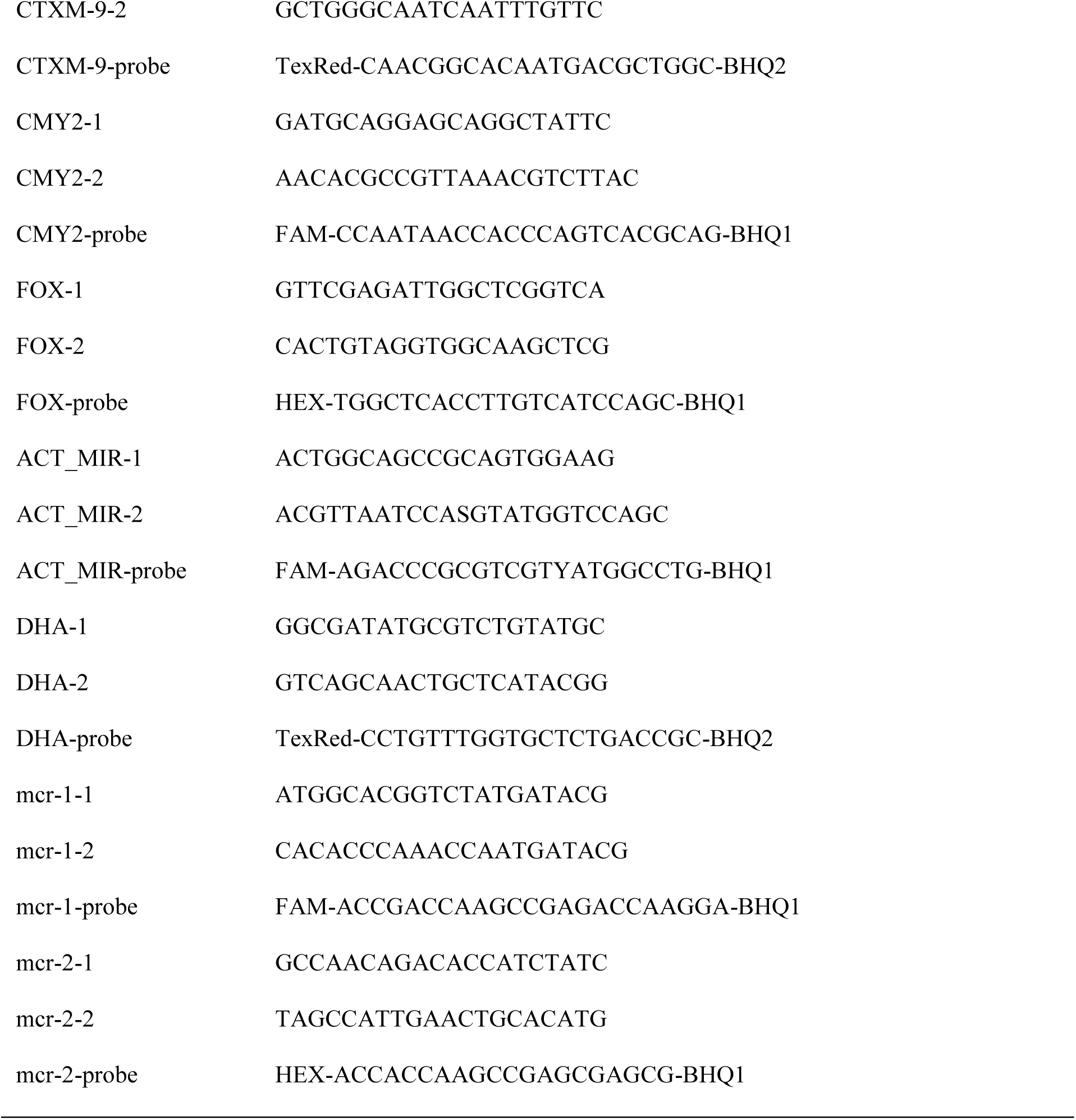
Primers used for the detection of genes encoding beta-lactamases (*bla*) and mobile colistin resistance (*mcr*).

**Table 2.**
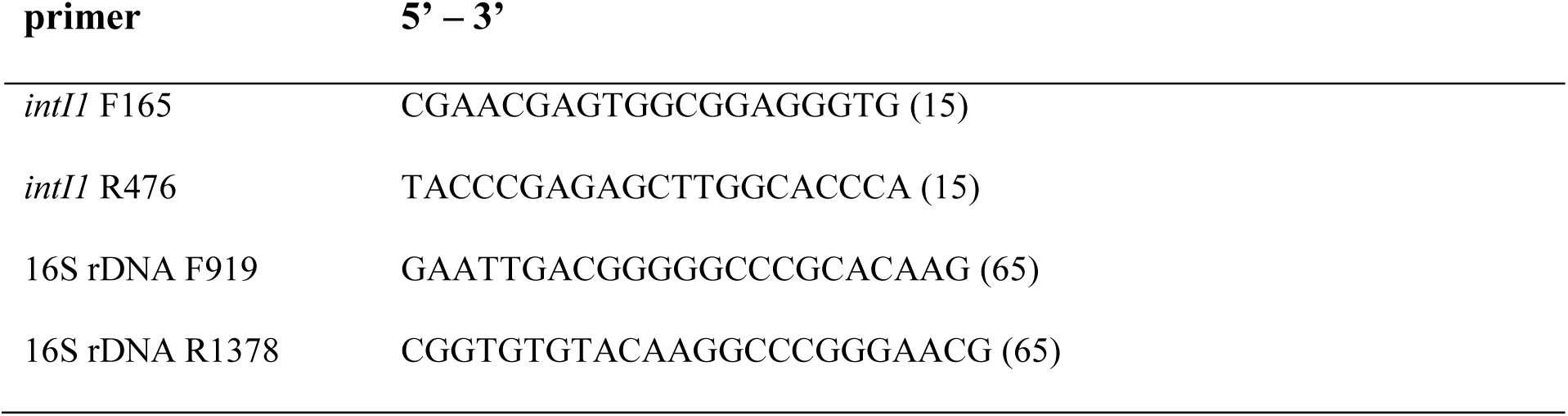
Primers used for the detection of class 1 integron-integrase gene (*intI1*) and 16S ribosomal DNA (16S rDNA).

### Isolation of antibiotic resistant bacteria

Samples were cultivated in tryptic soy broth (TSB; Merck, Darmstadt, Germany) containing subinhibitory concentrations of imipenem, cefotaxime or colistin to select resistant species. Subinhibitory concentrations were used since antibiotics normally do not occur in lethal concentrations in the investigated environments and in this way, the growth of susceptible strains should be inhibited while strains with only decreased susceptibility and resistances might still be isolated (32). In this respect, 5 mL of TSB containing 2 µg mL^-1^ imipenem or 6 µg mL^-1^ cefotaxime were inoculated with 100 µL of prepared samples, and 1 mL of TSB containing 10 µg mL^-1^ colistin were inoculated with 50 µL of prepared sample, respectively, to allow the isolation of carbapenemase- and ESBL-producing species as well as bacteria resistant to colistin. The inoculated broths were incubated for 24 h at 30 °C in an orbital shaker at 300 rpm. In case of growth, broth cultures were sub-cultivated on MacConkey agar (Carl Roth, Karlsruhe, Germany) with antibiotic-discs (24 h at 30 °C) of either imipenem, cefotaxime or colistin, depending on the antibiotic used in the broth, and on chromID agar for the isolation of carbapenemase-producing (chromID CarbaSmart, bioMérieux, Nürtingen, Germany) and ESBL-producing (chromID ESBL, bioMérieux) bacteria (24 h at 37 °C). After sub-cultivation on MacConkey agar, pure isolates were applied to VITEK 2 system for the determination of species and antimicrobial susceptibility testing (see below).

### Culture-based determination of species and antimicrobial susceptibility testing

After 24 h incubation on MacConkey agar at 30 °C, fresh isolates were applied to VITEK 2 compact system (bioMérieux) for the determination of bacterial species and resistance to antibiotics, according to the manufacturer’s instructions. Clinical resistance was assessed based on EUCAST breakpoints. VITEK 2 GN, VITEK 2 AST N-214 (ESBL-test, ampicillin, ampicillin/sulbactam, piperacillin/tazobactam, cefuroxime, cefuroxime/axetil, cefpodoxime, cefotaxime, ceftazidime, ertapenem, imipenem, meropenem, gentamicin, ciprofloxacin, moxifloxacin, tetracycline, tigecycline, trimethoprim/sulfamethoxazole) and VITEK 2 AST N-248 (piperacillin, piperacillin/tazobactam, cefotaxime, ceftazidime, cefepime, aztreonam, imipenem, meropenem, amikacin, gentamicin, tobramycin, ciprofloxacin, moxifloxacin, tigecycline, fosfomycin, colistin, trimethoprim/sulfamethoxazole) were used for the identification and analysis of antibiotic resistance of gram-negative bacteria, without prior determination of gram status. After determination of antibiotic resistances, DNA was extracted from identified isolates by a standard heat treatment (33) and qPCR was performed as well. Bacterial strains confirmed for producing ESBL by the VITEK 2 system were screened for *bla* genes belonging to the *bla*_CTX-M_, *bla*_CMY-2_, *bla*_FOX_, *bla*_ACT/MIR_ and *bla*_DHA_ families. Carbapenem-resistant isolates were screened for *bla*_VIM_, *bla*_NDM_, *bla*_KPC_, *bla*_GES_, *bla*_OXA-48_, *bla*_OXA-58_ and *bla*_OXA-23_ while colistin-resistant isolates were analyzed for *mcr-1* and *mcr-2* genes.

### Reduction of antibiotic resistant bacteria in washing machines and dishwasher

For investigating the antimicrobial effect of both washing machine and dishwasher, an antibiotic resistant strain and a susceptible strain of *Escherichia coli* (susceptible strain ATTC10536 and resistant strain isolated from a shower drain), *Klebsiella pneumoniae* (susceptible strain ATCC13883 and ESBL strain isolated from a wastewater treatment plant) and *Staphylococcus aureus* (susceptible strain ATTC6538 and MRSA strain CCUG35601) were used. Artificially contaminated cotton test swatches of 1 cm^2^ were prepared according to Honisch et al. (2014) for the analysis of the laundering process, while stainless steel biomonitors according to DIN EN 10088-3 were artificially contaminated and used for the dishwasher tests as described by Brands et al. (2020). To determine the effect of domestic laundering on antibiotic resistant strains, test runs were performed in a lab-scale washing machine (Rotawash) using the method of Schages et al. (2020). All tests were performed in triplicates; a 30 and 60 min main wash was tested at 30 °C and 40 °C using IEC-A* base powder (wfk testgewebe, Brüggen, Germany) as AOB-free detergent. Besides the main wash, the effect of the rinsing cycle was investigated with and without addition of quaternary ammonium compounds (QACs) using 0.02% benzalkonium chloride (BAC). The dishwasher tests were performed in an automated dishwasher (GSL-2, Miele & Cie. KG, Gütersloh, Germany) with each six repetitions and the following conditions were tested: five minutes main wash at 45 °C + rinsing cycle at 50 °C, 15 minutes main wash at 60 °C + rinsing cycle at 50 °C and 90 minutes main wash at 45 °C + rinsing cycle at 50 °C. After the tests, the logarithmic reduction (LR) was determined according to Block et al. (2001):

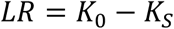

*K*_*0*_: initial load on the swatches/biomonitors before laundering/automated dishwashing

*K*_*S*_: remaining load on the swatches/biomonitors after laundering/automated dishwashing

### Statistics

Statistics were performed using GraphPad Prism (GraphPad Software Inc.). Data were expressed as means (± standard error). Data were log-transformed prior to statistical analyses to meet assumptions of normality and homogeneity of variance when needed. Absolute and relative abundance of *bla* genes, *intI1* genes and phenotypic resistance of bacteria in SD, WM and DW samples were normally distributed (Shapiro-Wilk test). Thus, statistically significant differences were assessed using two-way analysis of variance (ANOVA), and post-hoc Tukey’s multiple comparison test was performed to identify significant differences between the ARG groups of SD, WM and DW (*p* ≤ 0.05). Due to no Gaussian distribution, Spearman correlation (*p* ≤ 0.05) was performed to analyze the co-occurrence of ARGs and to determine the relation between phenotypic and genotypic resistance.

## Results

### Frequency distribution and co-occurrence of *bla* and *intI1* genes in households

In total, 207 beta-lactamase (*bla*) genes were identified in shower drains (SD, n=54), washing machines (WM, n=54) and dishwashers (DW, n=44), and 12 of the 13 targeted beta-lactamase types were detected while no mobile colistin resistance genes (*mcr-1* and *mcr-2*) occurred. *Bla*_VIM_, and *bla*_DHA_ only were identified in two SD samples and *bla*_CTX-M-9_ was detected once in a WM whereas the metallo-β-lactamase NDM occurred in none of the samples. The most frequently detected ARG was *bla*_CMY-2_ followed by *bla*_ACT/MIR_ and *bla*_OXA-48_. The absolute abundance of *intI1* and ampC-β-lactamase genes was significantly higher in SD samples compared to WM and of it was only significantly higher regarding ampC-β-lactamase genes compared to DW. In contrast, the relative abundance of ampC-β-lactamase genes, carbapenemase genes and total of ARGs was significantly higher in SD and DW vs. WM samples and of *intI1* and total ARGs in SD compared to DW samples while the abundance of carbapenemase genes was slightly higher in DW (Figure 1).

**Figure 1.**
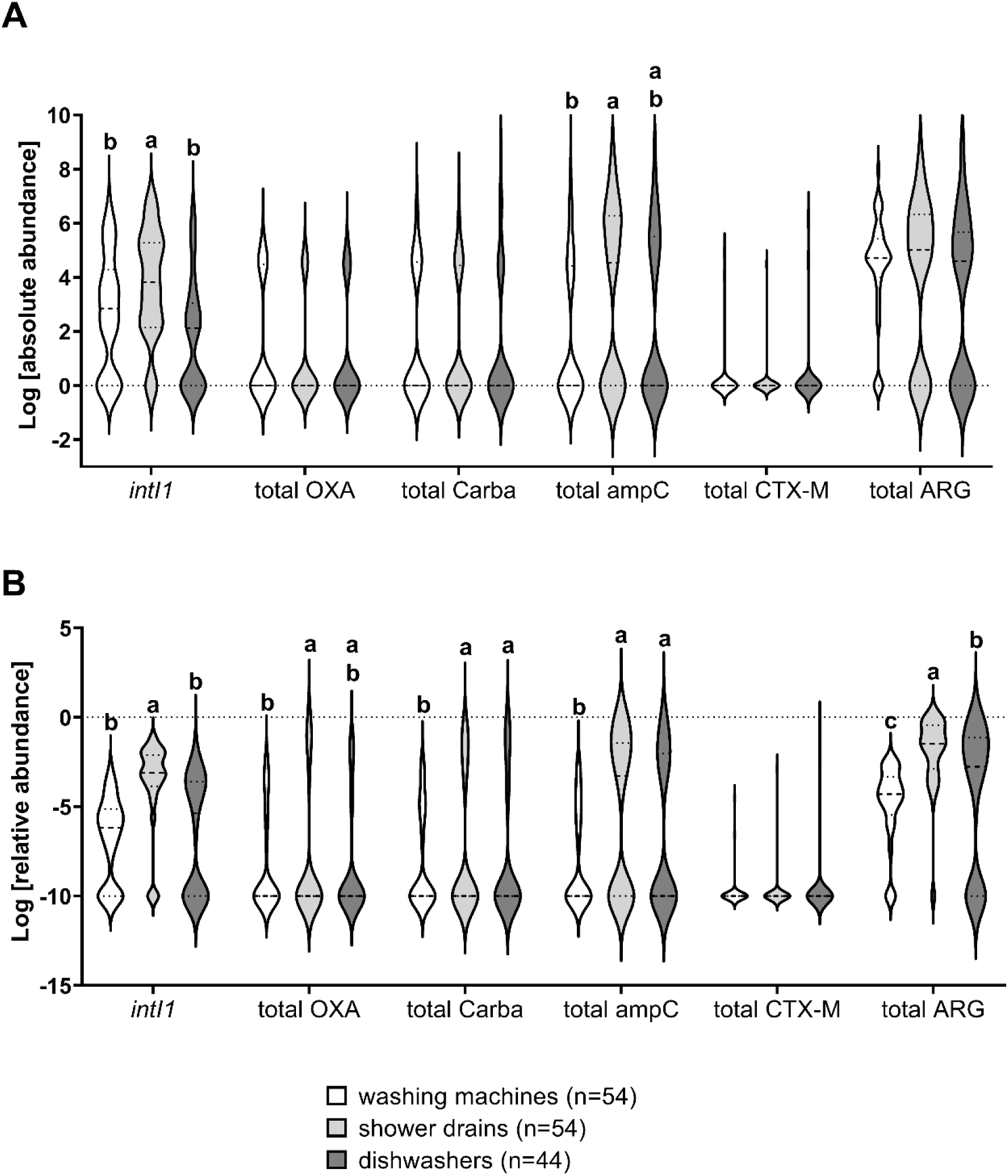
Absolute (A) and relative (B) abundance of *intI1* and *bla* genes in washing machines (WM), shower drains (SD) and dishwasher (DW) samples. The total of each gene group is presented as OXA: *bla*_OXA-58_ and *bla*_OXA-23_, Carba: *bla*_OXA-48_, *bla*_GES_, *bla*_KPC_ and *bla*_VIM_, ampC: *bla*_CMY-2_, *bla*_FOX_, *bla*_ACT/MIR_ and *bla*_DHA_ and CTX-M: *bla*_CTX-M-1_ and *bla*_CTX-M-9_. In B, samples without ARG (rel. abundance=0) are presented as -10. Different letters indicate significant differences at *p* ≤ 0.05 between SD, WM and DW of each group. Where no letters are shown, no significant differences were detected.

The number of detected ARGs in individual samples ranged from zero to five and the SD samples revealed the highest diversity with 24.1% containing three or more *bla* genes while in 40.9% of DW and 33.3% of WM samples no *bla* gene was detected (Table 3). Most ARGs were detected in SD (39.1%), followed by WM (33.3%) and DW (27.5%). An absolute abundance of ARGs of 1.53 × 10^7^ gene copies mL^-1^ in SD, 2.52 × 10^6^ gene copies mL^-1^ in WM and 2.74 × 10^7^ gene copies mL^-1^ in DW was determined. In contrast, the relative abundance was highest in SD samples (0.2367 ARG copies/16S rDNA copies) followed by DW (0.1329 ARG copies/16S rDNA copies) and WM (0.0006 ARG copies/16S rDNA copies).

**Table 3.** Co-occurrence of *bla* genes in shower drain (n=54), washing machine (n=54) and dishwasher (n=44) samples.

|  | shower drains | washing machines | dishwashers |
| --- | --- | --- | --- |
| **no *bla* gene** | 29.6% | **33.3%** | **40.9%** |
| **1 *bla* gene** | 24.1% | 31.5% | 20.5% |
| **2 *bla* genes** | 22.2% | 22.2% | 20.5% |
| **≥ 3 *bla* genes** | **24.1%** | 12.9% | 18.2% |

Co-occurrence of *bla* genes was explored using Spearman correlation, and the absolute abundance of total ARGs, *bla*_CMY-2_ and *bla*_ACT/MIR_ correlated strongly with *intI1* in SD and DW samples. Furthermore, *bla*_CMY-2_ and *bla*_ACT/MIR_ correlated strongly in SD and DW samples while inter alia *bla*_OXA-23_ and *bla*_OXA-48_ as well as *bla*_OXA-48_ and *bla*_FOX_ revealed a positive correlation (Figure 2). In DW samples, *bla*_OXA-58_ and *bla*_KPC_ correlated strongly whereas a moderate positive correlation of *bla*_GES_ and *bla*_CTX-M-1_ as well as *bla*_ACT/MIR_ and *bla*_DHA_ was observed in SD samples.

**Figure 2.**
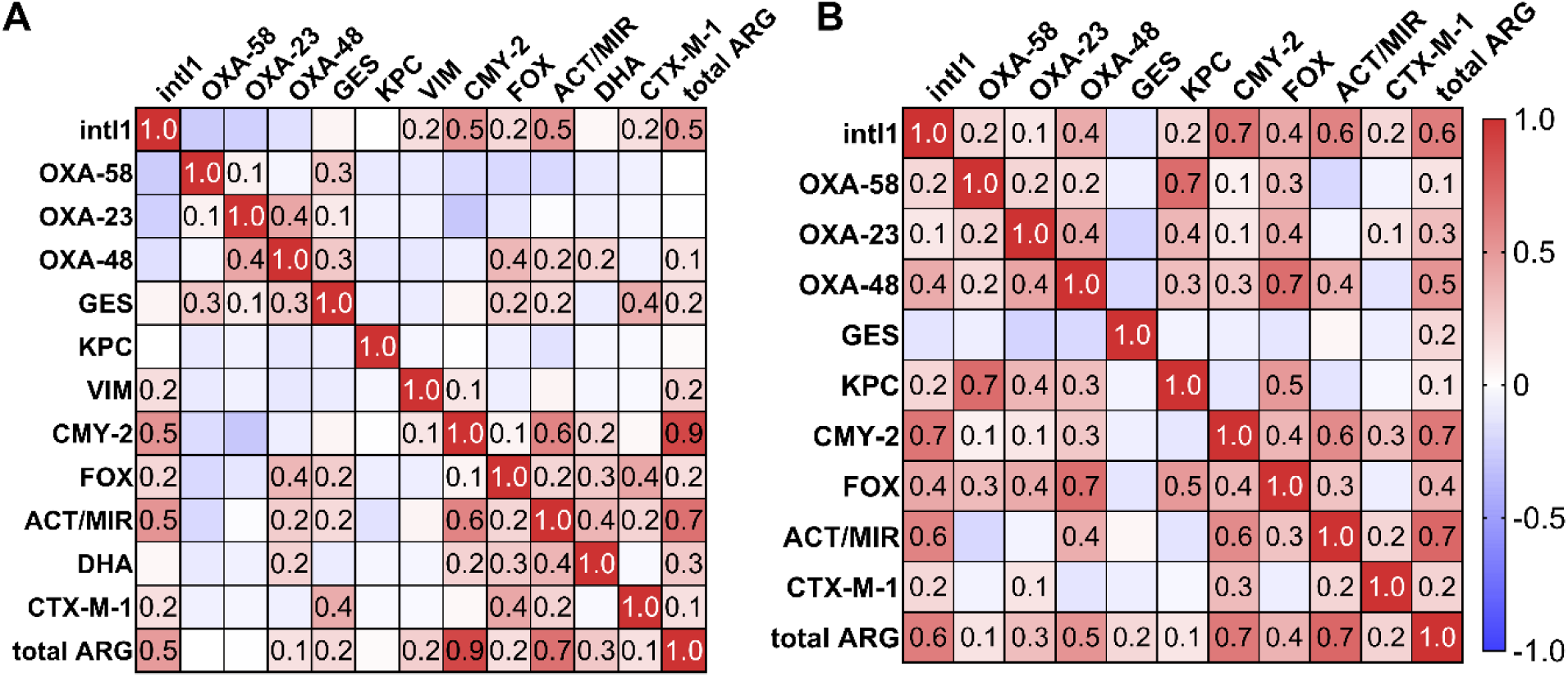
Correlation matrix showing co-occurrence of *bla* genes in shower drains (A) and dishwashers (B). Red indicates a strong positive correlation while blue indicates a strong negative correlation and a correlation of r ≥ 0.3 is statistically significant. *Bla* genes in washing machine samples did not correlate significantly.

### Abundance of ABR bacteria in household samples

A total of 273 strains was isolated from household samples using subinhibitory concentrations of antibiotics (Table 4). Intrinsic resistant species were predominant across household isolates (*p* ≤ 0.05, Tukey’s multiple comparison test), *Enterobacteriaceae* dominated in SD and DW and *Pseudomonas aeruginosa* was most abundant in WM (not significant). Except for the abundance of *Enterobacteriaceae* and *Pseudomonas* spp., bacterial species were distributed equally among SD, WM and DW. However, less strains were isolated from DW samples due to no detectable growth in supplemented TSB.

**Table 4.**
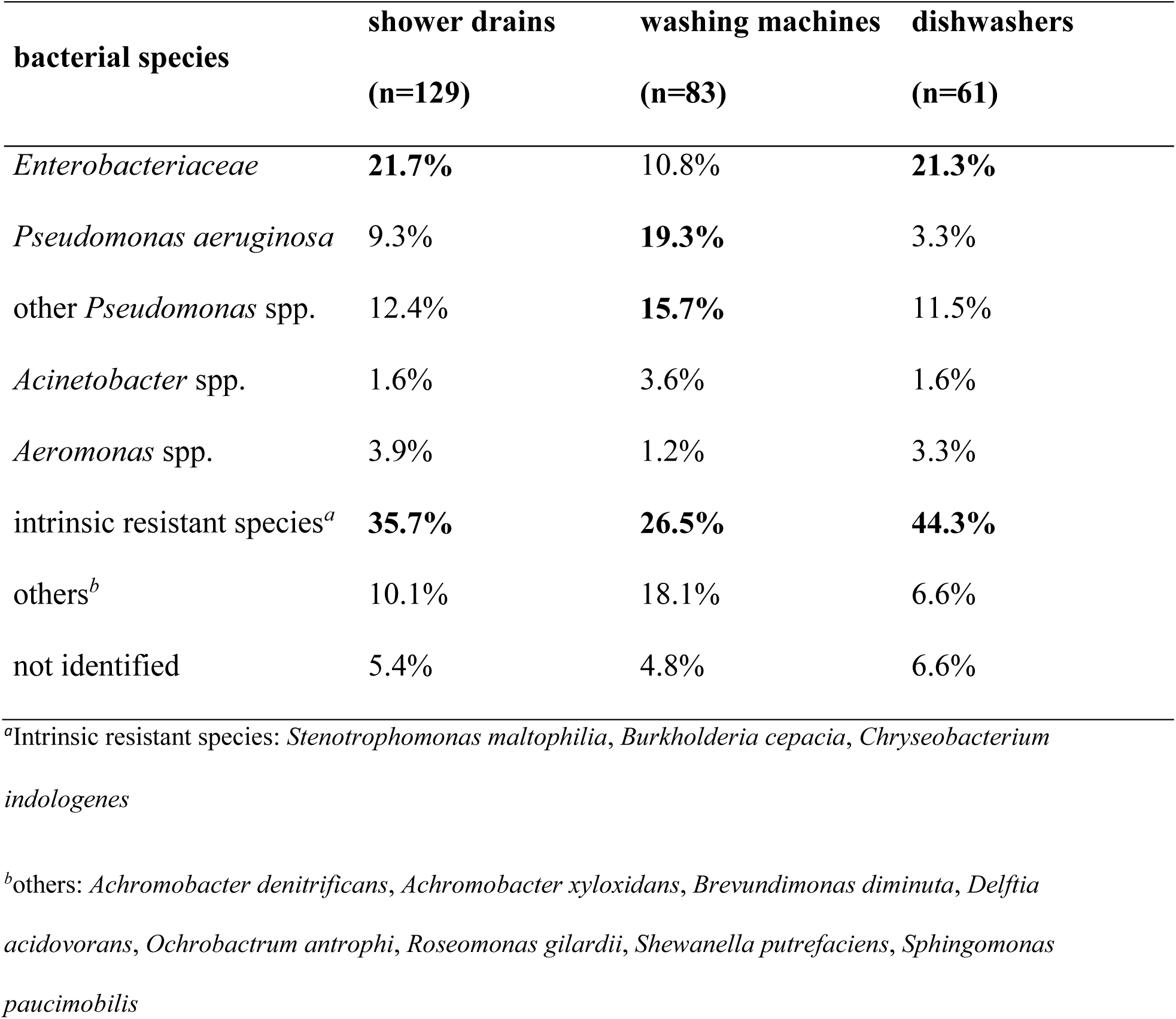
Percentage of resistant bacterial species isolated from shower drains, washing machines and dishwashers using subinhibitory concentrations of antibiotics.

Insusceptibility to carbapenems was significantly higher in SD samples, while in WM and DW intrinsic resistance dominated across the samples (Figure 3A). Resistance to cefotaxime/ceftazidime (CTX/CAZ) was higher in SD and DW samples compared to WM. Most of the identified species were environmental opportunistic pathogens, and although MDR and ESBL-producing bacteria were isolated from all samples without significant differences between SD, WM and DW (*p* ≤ 0.05), their contribution to resistant bacteria was highest in the SD samples.

**Figure 3.**
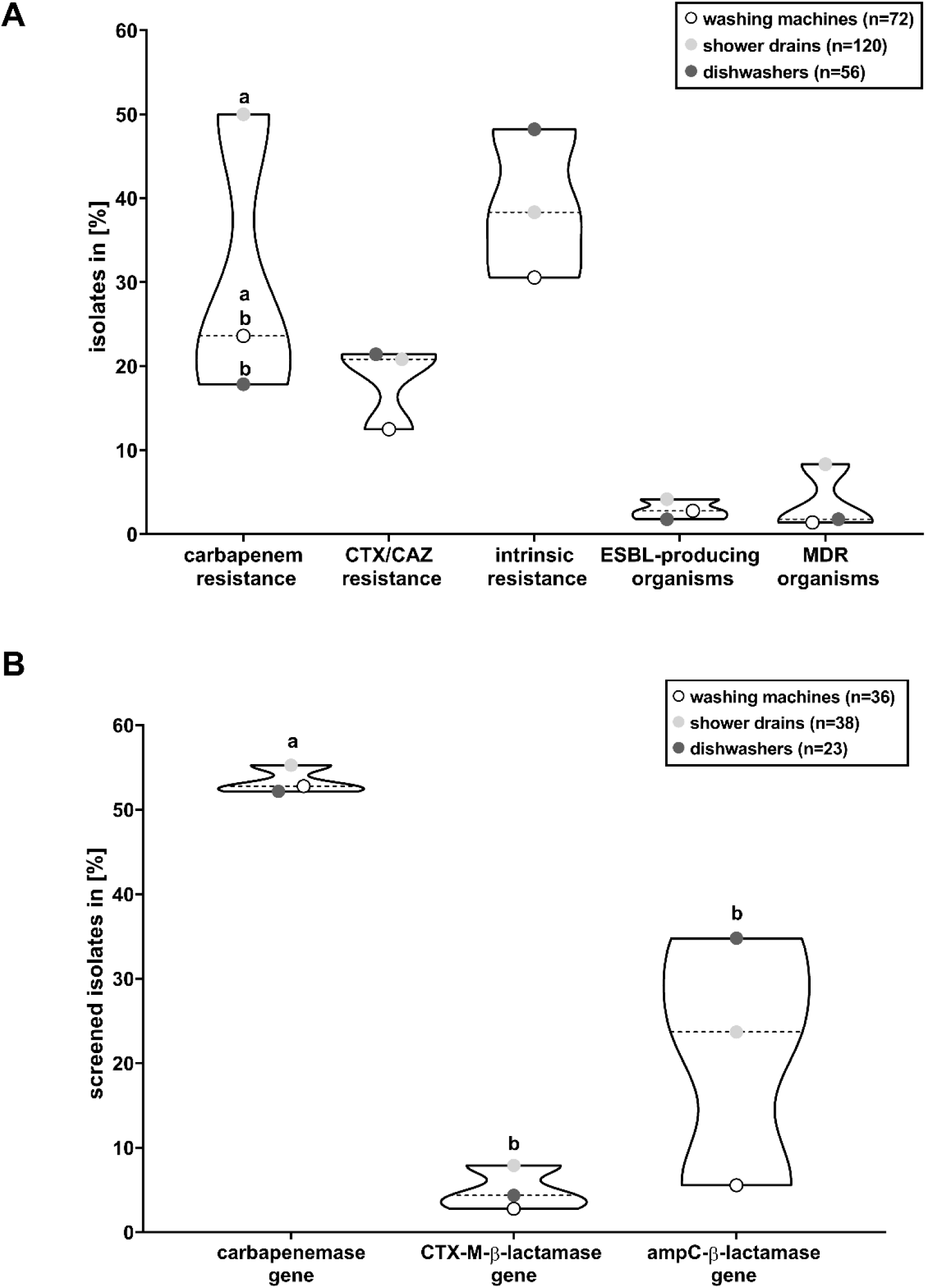
Percentage of resistant bacterial species (A) and *bla* genes detected in screened isolates (B) from shower drains (SD), washing machines (WM) and dishwashers (DW) of 54 different households. Resistances and ESBL-phenotype were determined using VITEK 2 system (bioMérieux), MDR bacteria were classified according to international recommendations and intrinsic resistance refers to the species *Stenotrophomonas maltophilia, Burkholderia cepacia* and *Chryseobacterium indologenes*. Different letters indicate significant differences at *p* ≤ 0.05 between SD, WM and DW of (A) and carbapenemase, CTX-M and ampC genes (B). Where no letters are shown, no significant differences were detected.

Quantitative PCR (qPCR) for the detection of *bla* and *mcr* genes was performed with the DNA extracted from isolates and in case of 33 of the 99 isolates the resistance profile of the phenotype was in accordance with the detected ARGs. Most *bla* genes were identified in strains of the *Enterobacteriaceae*, followed by *Pseudomonadaceae*. Carbapenemase genes were detected predominantly (*p* ≤ 0.05) while in approx. 50% of the isolates no ARGs were detected. Except for ampC genes, SD samples revealed the highest percentage of strains carrying ARGs (Figure 3B). Of the strains which were screened for *bla* and *mcr* genes, all MDR bacteria carrying ARGs were isolated from SD (see Table S1 and S2).

Spearman correlation of carbapenemase, CTX-M and ampC genes with phenotypic resistance against carbapenems, ceftazidime and/or cefotaxime (CAZ/CTX) and piperacillin/tazobactam (PIP/TAZ) was performed to identify the relevance of the detected ARGs (Figure 4). The high percentage of carbapenem resistant strains was confirmed by the detection of carbapenemase genes in more than 50% of the isolates and the strong positive correlation of carbapenemase genes and carbapenem resistance. PIP/TAZ and CAZ/CTX resistance correlated strongly with the ampC genes. Even though the occurrence of CTX-M genes correlated positively with CAZ/CTX resistance as well, the determined correlation was weak.

**Figure 4.**
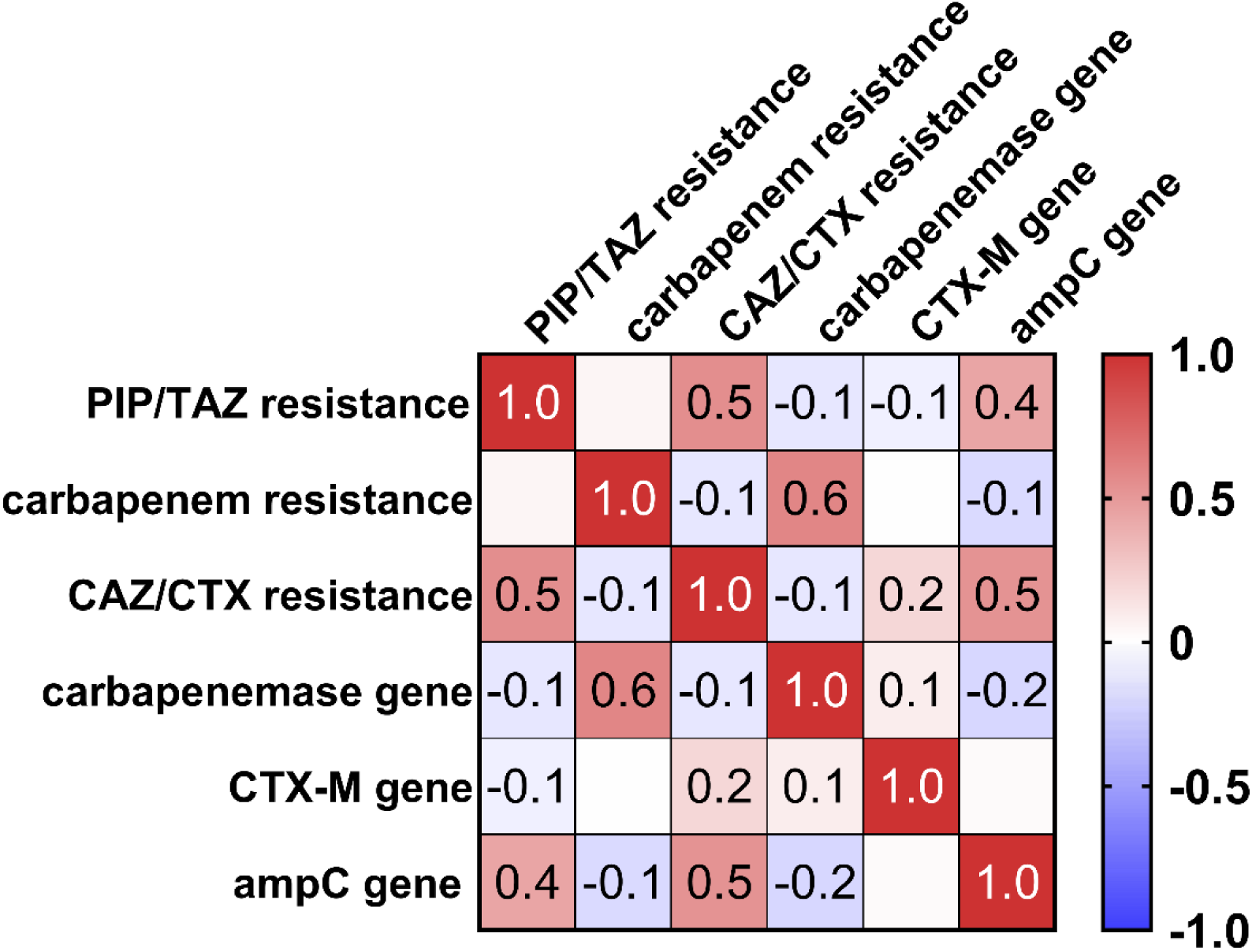
Spearman correlation (*p* ≤ 0.05) of genotype (carbapenemase genes: *bla*_OXA-58_, *bla*_OXA-23_, *bla*_OXA-48_, *bla*_GES_, *bla*_KPC_ and *bla*_VIM_, ampC genes: *bla*_CMY-2_, *bla*_FOX_, *bla*_ACT/MIR_ and *bla*_DHA_ and CTX-M genes: *bla*_CTX-M-1_ and *bla*_CTX-M-9_) with phenotypic resistance (to piperacillin/tazobactam (PIP/TAZ) carbapenems and ceftazidime and/or cefotaxime (CAZ/CTX)) in screened isolates from households.

### Effect of laundering and automated dishwashing on antibiotic resistant strains

To evaluate the persistence of ABR bacteria to laundry procedures compared to susceptible strains, artificially contaminated cotton swatches were washed in the Rotawash, simulating household relevant laundering parameters. An AOB-free detergent for the main wash and benzalkonium chloride (BAC) for the rinsing cycle were chosen, since the use of liquid or color detergents (without AOB) increases steadily in private households (38) and BAC is commonly used in rinse aids. Textile swatches were artificially contaminated resulting in initial microbial counts (expressed as colony-forming units (cfu) mL^-1^ of recovery fluid) of 2.72 × 10^7^ cfu mL^-1^ for *E. coli*, 3.29 × 10^7^ cfu mL^-1^ for carbapenemase-producing *E. coli*,c 2.25 × 10^7^ cfu mL^-1^ for *K. pneumoniae*, 3.01 × 10^7^ cfu mL^-1^ for ESBL-producing *K. pneumoniae*, 7.45 × 10^7^ cfu mL^-1^ for *S. aureus* and 3.38 × 10^7^ cfu mL^-1^ for MRSA. Although both susceptible and resistant strains were tested, no significant differences except for tests with BAC were revealed between the susceptible and ABR strain of the same species. Therefore, only results of the ABR strains are shown, and results of susceptible strains are provided in the supplemental material (see Figure S1).

Tests performed with AOB-free detergent in the Rotawash (Figure 5A) revealed only minor differences between the LR of carbapenemase-producing *E. coli*, ESBL-producing *K. pneumoniae* and MRSA. In general, the results indicate an increasing efficacy of laundering with increasing time and temperature, revealing significantly higher reductions of all strains at 40 °C/60 min compared to tests at 30 °C. The tests of the rinsing cycle showed that the effect of 0.02% BAC seemed to be negligible, except for the significantly higher reduction of the susceptible *E. coli* strain (Figure 5B). However, the highest reductions achieved were in the range of 45 to 57% at 40 °C and a main wash of 60 min.

**Figure 5.**
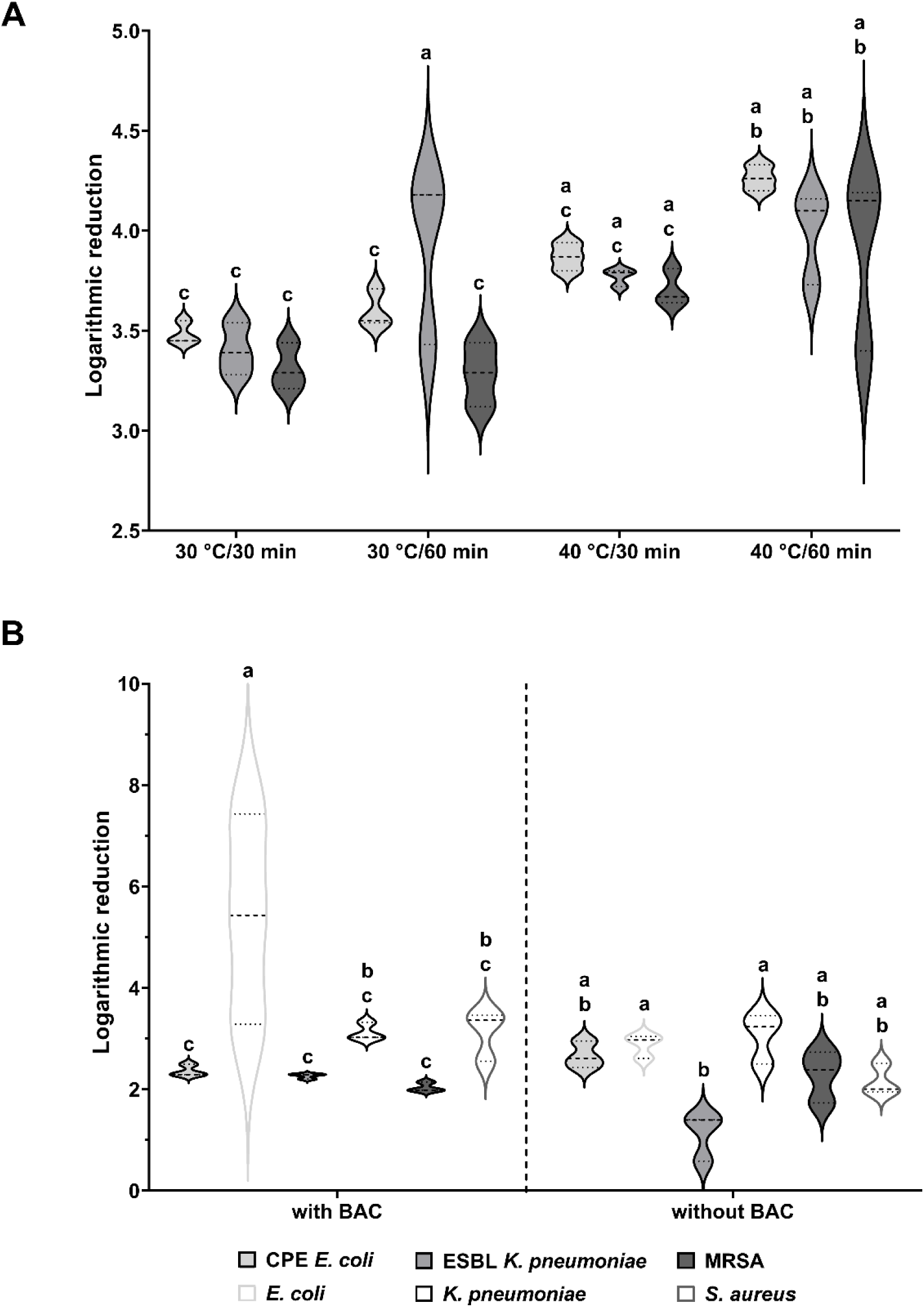
Impact of main wash with AOB-free detergent on ABR strains (A) and rinsing cycle with and without benzalkonium chloride (BAC) (B) of the laundering process simulated using the Rotawash on a resistant and susceptible strain of *E. coli, K. pneumoniae* and *S. aureus*. Different letters indicate significant differences at *p* ≤ 0.05.

The same test strains were used to determine the effect of automated dishwashing on ABR strains. The initial microbial counts of the contaminated biomonitors were 6.82 × 10^7^ cfu mL^-1^ for *E. coli*, 7.18 × 10^7^ cfu mL^-1^ for carbapenemase-producing *E. coli*, 1.40 × 10^7^ cfu mL^-1^ for *K. pneumoniae*, 1.20 × 10^8^ cfu mL^-1^ for ESBL-producing *K. pneumoniae*, 1.55 × 10^8^ cfu mL^-1^ for *S. aureus* and 1.62 × 10^8^ cfu mL^-1^ for MRSA. In order to simulate household relevant conditions, the following programs were tested using reference detergent: five min main wash at 45 °C (short program), 15 min main wash at 60 °C (standard program) and 90 min main wash at 45 °C (eco-program). Again, only the ABR strains are shown since the results between ABR (Figure 6) and susceptible strains (Figure S2) did not differ significantly.

**Figure 6.**
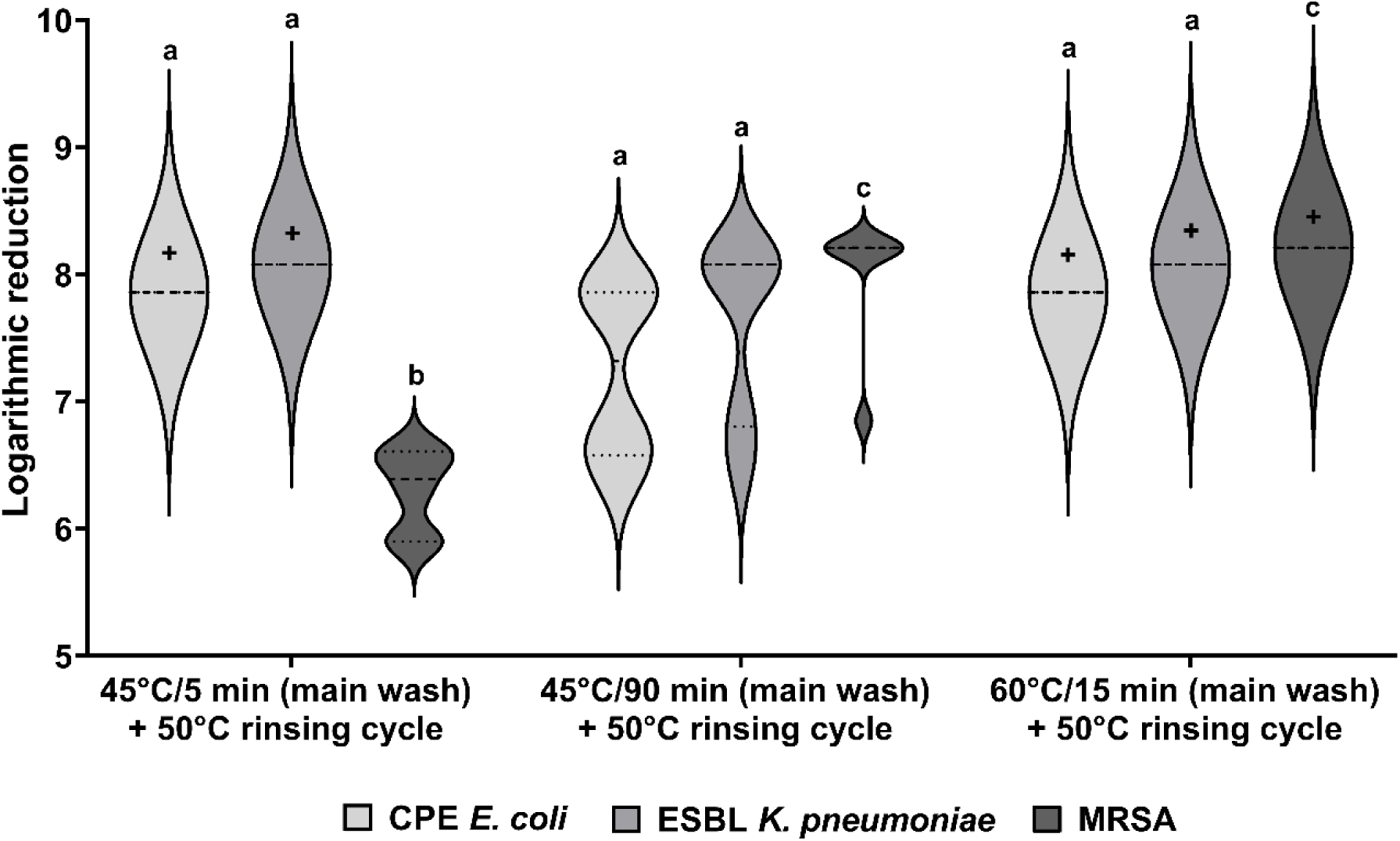
Impact of automated dishwashing on ABR strains of *E. coli, K. pneumoniae* and *S. aureus* with detergent. The different values for LR max. [+] (indicating a complete reduction of the microbial load in case of 45 °C/ min and 60 °C/15 min) were obtained due to different initial loads on the biomonitors. Different letters indicate significant differences at *p* ≤ 0.05.

Regarding all ABR strains, a complete reduction was achieved at 60 °C, and only the LR of MRSA was significantly lower at both 45 °C programs (Figure 6). Even under the conditions of the short program the LR max. (exceeding the detection limit) was reached except for MRSA while the eco-program revealed lower LR. But with all programs the reductions were at least above 75%.

## Discussion

Previous studies revealed the presence of *bla* genes (26) and the correlation of multi-resistance with *intI1* (16) in the domestic environment, but no studies on the frequency distribution of *bla* genes and their co-occurrence in private households, especially in relation to phenotypic resistance, have been conducted so far. SD, WM and DW provide a good habitat for microbial communities with a steady nutrient supply in a humid environment, promoting bacterial growth and biofilm formation (39–42). Besides, ABR bacteria and antibiotic residues have been detected in siphons of hospitals (43, 44), indicating their occurrence in SD of domestic areas as well. In this study, we detected *bla* genes in SD (70.4%), WM (66.7%) and DW (59.1%) and isolated ABR bacteria from 81.5% of all households. Bacterial isolates carrying multiple ARGs were identified and SD revealed the highest prevalence of both ARGs and ABR bacteria. Laundering led to significant reductions of ABR bacteria, although bacterial counts of approx. 10^3^ cfu mL^-1^ were still detected after the tests. Automated dishwashing achieved higher logarithmic reductions (LR) and in most cases no bacterial growth was detected afterwards.

### Prevalence and co-occurrence of ARGs and ABR bacteria in household samples

The obtained results show a strong variation of all *bla* genes across SD, WM and DW samples due to very high absolute abundances of ARGs (*e*.*g*. 4.04 × 10^8^ copies mL^-1^ of carbapenemase genes in a DW sample) compared to a high amount of negative results (*e*.*g*. 95.5% of all household samples in case of CTX-M genes). However, relative abundance of ARGs was higher in SD samples except for CTX-M genes, indicating a higher frequency of ARGs in SD. AmpC-β-lactamase genes occurred predominantly in all samples and dominated in a previous study of WM and DW as well (26).

Correlation analysis revealed significantly positive co-occurrence of *bla* genes such as *bla*_CMY-2_ and *bla*_ACT/MIR,_ *bla*_OXA-23_ and *bla*_OXA-48_ or *bla*_GES_ and *bla*_CTX-M-1_ in SD and DW. Since those ARGs occur plasmid-mediated (4), a transfer of the genes and thus a high prevalence resulting in co-occurrence within samples seems likely. Furthermore, in the majority of SD, WM and DW samples and in 43.9% of isolates carrying *bla* genes more than one *bla* gene was detected. This is supported by Ju et al. (2016) revealing co-occurrence of ARGs that shared the same resistance mechanism since the selective pressure of the same antibiotic results in the co-selection of the same ARG types. Studies revealed that gene cassettes within class 1 integrons carry carbapenemase, ESBL-, ampC- and OXA-β-lactamase genes (46–48), indicating that the detected *bla* genes in this study might be partially located on class 1 integrons as well, since total ARG abundance and *intI1* correlated strongly.

The bacterial community of SD, WM and DW usually comprises environmental bacteria, including, among others *Enterobacteriaceae* and *Pseudomonadaceae* (26, 39, 41), which were the predominant species in this study as well apart from intrinsic resistant species such as *Stenotrophomonas maltophilia*. The identified species are known to harbor *bla* genes (4, 10, 49), and the isolation of clinically relevant species such as MDR *P. aeruginosa* and ESBL-*E. coli* harboring carbapenemase genes suggests that the domestic environment might be a potential reservoir for the spread of beta-lactam resistance. The positive correlation of carbapenemase and ampC-β-lactamase genes with carbapenem resistance and cefotaxime/ceftazidime, respectively, shows that the phenotypic resistance is most likely based on beta-lactamase production. Interestingly, in 21 out of 54 households at least one detected *bla* gene was the same in all samples (12 households), in SD and DW (five households), in SD and WM (four households) or in WM and DW (two households). The connection of the sample points to the water supply system and the growth of biofilms harboring diverse microbial communities on inner pipe surfaces (39, 50) might contribute to the spread of ARGs and ABR bacteria from/to and within households. In addition to the water supply system, a transfer within the same household by household members or contaminated eating utensils/laundry is likely as well (51–53) and has been shown by Schmithausen et al. (2019) in a WM.

### Shower drains as a hotspot of ARGs and ABR in private households

The significantly higher relative abundance (*intI1*, total of carbapenemase, OXA-β-lactamase and ampC-β-lactamase genes) in SD samples compared to WM and partially to DW samples might indicate a higher resistance potential. Shower drains are known to be prone to biofilm formation, which are more resistant to environmental factors (such as exposure to antibiotics), promote horizontal gene transfer and thus ABR (54–56). Therefore, the high abundance of ARGs, especially of ampC-β-lactamase genes (50), might be related to biofilms. Domestic drains are colonized with *Pseudomonadaceae* and coliform bacteria (41), which dominated in the investigated SD samples and are associated with beta-lactamases, multi-drug resistance and nosocomial outbreaks originating from contaminated sinks (57, 58). We found that strains harboring *bla* genes, ESBL-producing and MDR bacteria dominated in SD, which indicates a higher resistance potential compared to WM and DW as well. This is supported by the identification of shower and sink drains as hotspots for *intI1* compared to other household areas by Lucassen et al. (2019) and by a study of the frequency of ABR bacteria in households of Marshall et al. (2012), revealing highest titers of ABR bacteria in sink drains. Thus, dissemination of ABR bacteria colonizing the human body to SD and their selective accumulation due to the exposure to biocides used in cleaning agents and personal care products (27, 59) seems likely.

### ABR bacteria in washing machines and dishwashers

The analysis of the impact of laundering and automated dishwashing on ABR bacteria showed that the efficacy increased with increasing duration and temperature. The reduction rates of resistant strains and non-resistant strains in both WM and DW were quite similar, and only MRSA and *S. aureus* revealed higher persistence, which was determined of both resistant and non-resistant strains in other studies as well (26, 60). However, to achieve higher reductions during laundering, higher temperatures or the use of a heavy-duty detergent containing a bleaching agent might be necessary (38) since bacterial counts of approx. 10^3^ cfu mL^-1^were detected even after laundering at 40 °C for 60 min. Thus, laundering at lower temperatures when using bleach-free detergents might lead to cross-contaminations of the laundry or the WM (29, 61) with ABR bacteria, which is especially of great concern regarding the rising number of elderly and ill people being nursed at home. Besides, no significantly higher reduction of ABR bacteria was observed when using 0.02% BAC in the rinsing cycle, questioning the need of such rinse aids since the use of quaternary ammonium compounds is suspected to co-select for ABR (62). In dishwashers, no bacterial growth was detected after test runs except for the eco-program, thus cleaned dishes seem to be no source of the tested strains. However, ABR bacteria were isolated from rubber door seals and DW sieves and have already been identified as a source of pathogenic and opportunistic pathogenic bacteria and fungi (42, 63, 64). Zupančič et al. (2019) determined a low phenotypic ABR potential in DW, which is confirmed by our study since less resistant strains were isolated compared to SD and WM. Nevertheless, DW samples revealed a high absolute abundance of ARGs, and the isolated strains still harbored *bla* genes and thus can be considered as a potential source of ABR bacteria as well.

## Conclusion

The results obtained in this study substantiate that the domestic environment represents a potential reservoir of *bla* genes and beta-lactam resistant bacteria. We found that ARGs co-occurred in household samples, indicating that bacterial species harboring multiple *bla* genes or bacterial communities harboring multiple beta-lactam resistant species are frequent in SD, WM and DW. This evidence is supported by the detection of various *bla* genes in the bacterial isolates. Moreover, our data show that SD seem to be the dominant source of beta-lactam resistance in households, revealing a higher frequency of *bla* genes, MDR and ESBL-producing bacteria. Although laundering and automated dishwashing significantly reduced ABR bacteria, low bacterial counts were still detected especially after laundering. Thus, a transfer of ABR bacteria via contaminated laundry or the grey water of WM in other environments seems to be possible. Further studies of the domestic environment are needed to confirm the possible risk of dissemination of ARGs and ABR bacteria and to determine whether households contribute to the spread of ABR or are a habitat where resistant bacteria from the environment, humans, food or water accumulate.

## Acknowledgements

This research was funded by the German Federal Environmental Foundation (Deutsche Bundesstiftung Umwelt) in a project analyzing potential transfer pathways of antibiotic resistance between the environment and households (Az 34632/01) and by the HSRW-scholarship for PhD students.

